# Multiscale Invasion Assay for Probing Macrophage Response to Bacteria

**DOI:** 10.1101/2020.11.16.385617

**Authors:** Kimberly A. Wodzanowski, April M. Kloxin, Catherine L. Grimes

**Affiliations:** Department of Chemistry and Biochemistry, University of Delaware, Newark, DE 19716; Department of Chemical and Biomolecular Engineering, University of Delaware, Newark, DE 19716; Department of Materials Science and Engineering, University of Delaware, Newark, DE 19716; Department of Biological Sciences, University of Delaware, Newark, DE 19716

## Abstract

The immune system is a complex network of various cellular components that must differentiate between pathogenic bacteria and the commensal bacteria of the human microbiome, where misrecognition is linked to inflammatory disorders. Fragments of bacterial cell wall peptidoglycan bind to pattern recognition receptors within macrophages, leading to immune activation. To study this complex process, an approach for three-dimensional (3D) culture of human macrophages and their invasion with relevant bacteria in a well-defined hydrogel-based synthetic matrix inspired by the gut was established. Workflows were developed for monocyte encapsulation and differentiation into macrophages in 3D culture with high viability. Bacteria invaded into macrophages permitted *in situ* peptidoglycan labeling. Macrophages exhibited biologically-relevant cytokine release in response to bacteria. This multi-dimensional bacteria-macrophage co-culture system will prove useful in future studies to observe bacterial fragment production and localization in the cell at the carbohydrate level for insights into how our immune system properly senses bacteria.

**TOC Figure:** 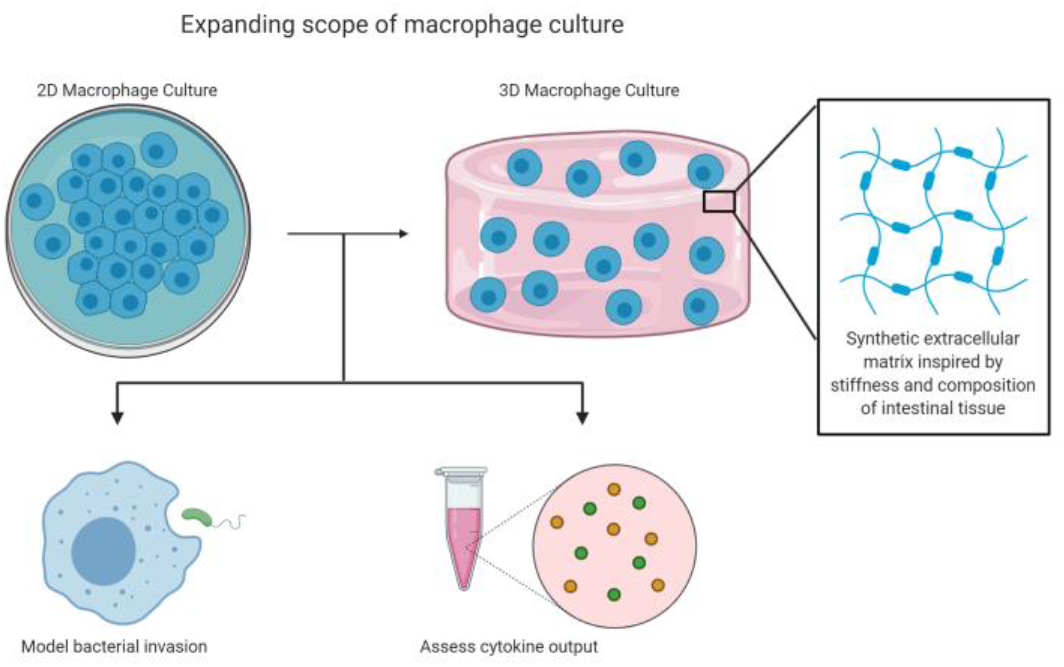

## Introduction

The innate immune system is the body’s first line of defense against invading pathogens and has developed molecular mechanisms to sense and differentiate between pathogenic bacteria and the over 39 trillion bacteria constituting the human microbiome. Immune cells called macrophages have many roles in the innate immune system, including ingesting pathogens by phagocytosis, scavenging dead cells and cell debris, and remodeling tissues after injury.^1^ Macrophages are derived from monocytes, which circulate in the blood stream until entering the tissue and differentiating into macrophages, based on release of activating lymphokines from T lymphocytes present in the area of infection.^2^ Mature macrophages express receptors that identify pathogens, allowing their proper uptake into the cell for degradation and response.^1^ These receptors include membrane-associated Toll-like receptors (TLRs) and cytosolic NOD-like receptors (NLRs), which are known to bind to fragments of the bacterial peptidoglycan (PG), a component of the bacterial cell wall. Chemists have developed synthetic PG mimics of smaller fragments such as muramyl dipeptide (MDP) and muramyl tripeptide (MTP), which bind to NLRs and therefore have been used to study immune responses (Figure 1). Misrecognition of various PG fragments by the immune system is hypothesized to lead to diseases including Crohn’s disease, inflammatory bowel disease, asthma, and gastrointestinal cancers.^3,4^ Although MDP and MTP serve as important tools in studying immune responses in humans, the true identity of naturally produced immunostimulatory fragments, how they are generated, and how they interact with innate immune receptors in macrophages is not well known.^5–7^ Therefore, there is a need for development of co-culture systems to model invasion of bacteria within macrophages to begin probing these complex questions.

**Figure 1:**
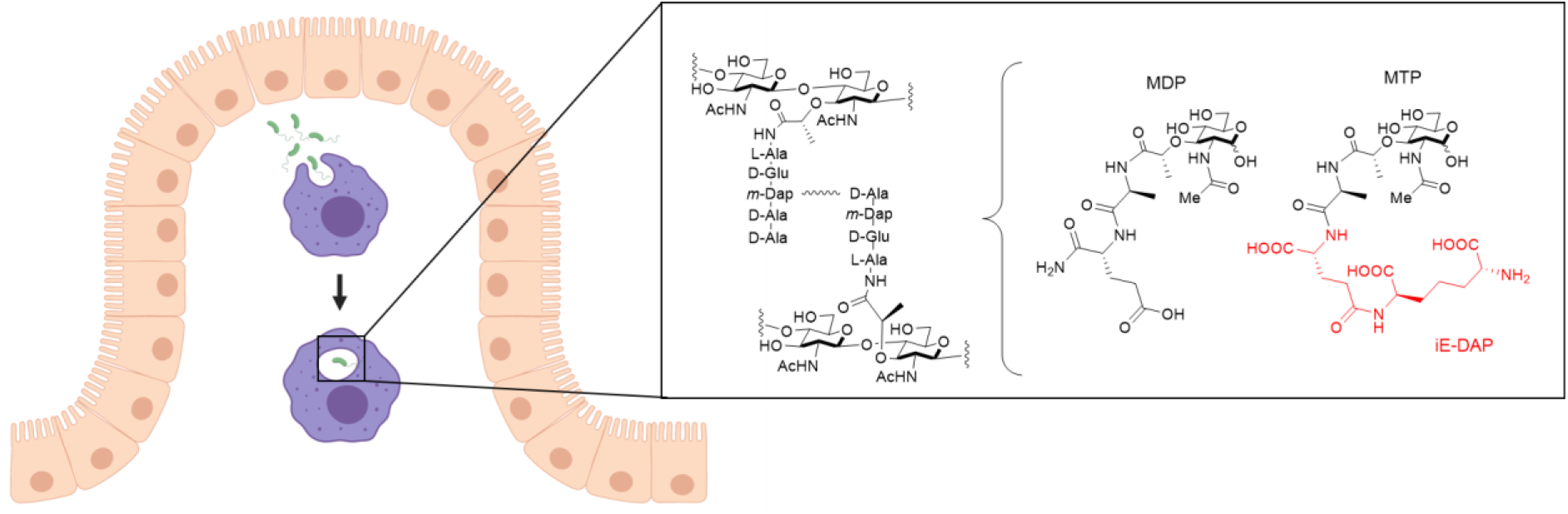
Invasion of bacteria into macrophages. Macrophages (purple) that arise from monocytes reside in the connective tissue underlying the intestinal epithelium (beige). Macrophages phagocytose pathogens such as bacteria (green) and break them down into smaller fragments to determine what type of signaling for an immune response is needed. One recognition element of bacteria, peptidoglycan (PG), is composed of alternating *N*-acetyl glucosamine (NAG) and *N*-acetyl muramic acid (NAM) sugars with a pentapeptide chain off the 3-position of the NAM that is further crosslinked to form mature PG. Out of this larger structure, smaller synthetic fragment, smuramyl dipeptide (MDP) and muramyl tripeptide (MTP), can be teased out with the core NAM sugar, which are known to elicit an immune response. D-glutamyl-meso-diaminopimelic acid (iE-DAP), shown in red, is a specific muropeptide in Gram-negative bacteria that activates the NLR, NOD1, to initiate an immune response.

Macrophages for studies of bacteria cell invasion are typically cultured on two-dimensional (2D) surfaces such as tissue culture polystyrene (TCPS). While this approach provides a well-defined environment, the material is unnaturally polarizing and has mechanical properties (Young’s modulus (E) ~ 3 GPa) a million times stiffer than those of the native soft tissue of the intestines (E ~ 2-20 kPa).^8,9^ Seminal works have demonstrated how the function of many different cell types (e.g., stem cells, epithelial cells, cancer cells) is influenced by the dimensionality and stiffness of the culture environment.^10–12^ Soft three-dimensional (3D) culture systems have been shown to be particularly effective for probing cell differentiation and migration in a physiologically relevant manner.^13^ As macrophages reside within connective tissue underneath the epithelium in the gut and respond to bacteria that breach this layer, 3D migration and interactions with the extracellular matrix (ECM) play an important role in infection clearing, suggesting the potential importance of studying macrophage response to bacterial invasion in multi-dimensional systems.^14^

For studies of the gut in three dimensions, recent progress has been made in the development of organoids cultures, amongst other approaches.^14,15^ Organoids are 3D cell clusters formed *in vitro*, with stem cells or cells derived from primary tissues, and are capable of self-organization and self-renewal, exhibiting similar function to *in vivo* organs.^16^ However, organoids lack immune cells, in addition to requiring lengthy timescales with intensive maintenance to fully generate (e.g., 1-3 months).^16,17^ Animal models have also provided insights in the context of inflammatory bowel disorders (IBD) by allowing study of mucosal inflammation. However, these models can have reproducibility issues and do not sufficiently capture human IBD as they cannot accurately control for intestinal pathology, inflammation, and bacteria related to IBD.^14,18–20^ Currently, a gap remains in physiologically-relevant, multi-dimensional systems for studying specific bacteria-immune cell interactions, where few immune cells, particularly macrophages, have been cultured in three dimensions.^21–24^ There is a need for robust hybrid systems with well-defined 3D properties that can be manipulated to reflect aspects of the native tissue and are suitable for co-culture of macrophages and bacteria.

Hydrogels, crosslinked water-swollen networks of hydrophilic polymers, have emerged as good candidates for mimicking a variety of soft tissue microenvironments for 3D cell culture applications. In particular, synthetic hydrogels can be engineered to mimic key properties of the native ECM, including mechanical properties and biochemical content, and permit 3D cell encapsulation and culture.^25^ These systems include multi-arm polymers (e.g., biologically-inert poly(ethylene glycol) (PEG)) that have been functionalized with reactive handles (e.g., (meth)acrylates, norbornenes, vinyl sulfones, thiols) for crosslinking with functionalized cell-degradable and integrin-binding peptides inspired by native tissues for a range of applications.^25^ To date, studies of macrophages with such synthetic materials largely have focused on probing the foreign body response in 2D culture studies^26,27^ with recent efforts demonstrating feasibility of macrophage 3D culture.^21,28^ These versatile materials provide significant opportunities for creating multi-dimensional culture systems with well-defined and tunable properties for probing macrophage-bacteria interactions to test hypotheses in more physiologically-relevant environments.

In this work, we aimed to establish a well-defined, bio-inspired co-culture system that enabled encapsulation and culture of immune cells relevant to the human gut, and to study their response to invasion by different types of Gram-negative bacteria in three dimensions. First, we identified a PEG-peptide hydrogel composition for achieving relevant synthetic matrix properties inspired by the healthy gut. Next, we established workflows for the successful encapsulation of human monocytes and their differentiation into human macrophages within these hydrogel-based synthetic matrices, inspired by the natural progression of immune cell infiltration and differentiation within native gut tissues. Following bacterial invasion, we bioorthogonally labeled bacterial PG and utilized these fluorescently labeled species to visualize macrophage engulfment of bacteria in both 2D and 3D culture. Further, we probed macrophage cytokine expression in these systems with contrasting dimensionality, elucidating key differences in macrophage response.

## Experimental

See supplementary information (SI) for synthetic and cell culture methods.

### Cell Encapsulation

THP-1 cells were encapsulated as a single cell suspension at a density of 5×10^6^ cells per mL in 20 μL of hydrogel precursor solution (250,000 cells/hydrogel). Precursor solution was prepared using 7mM PEG-8-Nb, 5mM linker peptide, 2mM pendant peptide, 2mM LAP, and THP-1 cell suspension in PBS, with all concentrations for the functional handle. Hydrogels were formed in 10 mm x 0.5 mm sterile gasket molds (43 μL precursor solution/mold) upon irradiation with a cytocompatible dose of long wavelength UV light (10 mW/cm^2^, 365 nm, 2 min; Omnicure 2000 with light guide and collimating lens). Two replicates were formed at a time and then placed in a 24 well plate with 500 μL of RPMI media. 200nM PMA was added to the media in each well for differentiation into macrophages. Cell-hydrogel constructs were incubated under sterile conditions at 37°C with 5% CO_2_.

### Live/Dead viability assay on encapsulated cells

Cell viability in the hydrogels was assessed using a LIVE/DEAD^®^ Viability/Cytotoxicity Kit (ThermoFisher Scientific) and imaged using confocal microscopy, which are described in SI.

### Flow cytometry

To confirm differentiation of THP-1 cells into macrophages within 3D culture, hydrogels with encapsulated THP-1 cells treated with PMA (for differentiation into macrophages) or untreated THP-1 monocytes were washed twice in 2 mL in 1×PBS for 5 min each. Hydrogels were put into 1.5 mL Eppendorf tubes (4 hydrogels per tube), and a 1mL solution of collagenase (300 U/mL) was added to degrade the hydrogel. Tubes were placed in CO_2_ incubator at 37°C. The solution was triturated every 10 min for up to 30 min until the solution could be pipetted freely. The now-digested hydrogel solution was centrifuged (150 g, 5 min) to pellet the cells. Monocytes (THP-1 cells) or macrophages (differentiated THP-1 cells) from 2D culture (1×10^6^ cells) were removed from plates and similarly centrifuged in 1.5 mL Eppendorf tubes. Pelleted cell samples were washed with 2% bovine serum albumin (BSA) in PBS and resuspended in 100 μL of 2% BSA in PBS, and 5 μL of CD11b antibody was added. Samples were placed on ice in the dark for 30 min and then centrifuged followed by washing with 2% BSA in PBS. Cells were then fixed in 4% paraformaldehyde in 1xPBS for 15 min at room temperature, centrifuged, and washed 2×100μL in Intracellular Staining Permeabilization Wash Buffer (BioLegend). Cells were resuspended in 100μL of Intracellular Staining Permeabilization Wash Buffer, and 5 μL of CD68 was added to each sample followed by incubation on ice for 30 min. Cells were washed 1×100 μL of Intracellular Staining Permeabilization Wash Buffer. Flow cytometry was performed on ACEA Novocyte Flow Cytometer. Samples were briefly vortexed before each run. 100,000 cell counts were collected for each sample and were analyzed in triplicate, and fluorescence intensities (height) were generated and overlaid.

### Invasion assay in 2D culture for Imaging

Sterile cover glasses (Fisher Scientific, catalogue number 12-545-80) were coated with 500 μL of 0.1 mg/mL poly-L-ornithine (Sigma-Aldrich) in 24 well plate overnight. The poly-L-ornithine was removed, and the cover glasses were washed with PBS twice. THP-1 cells were seeded on the cover glasses in 24-well plates with RPMI media (1 x 10 ^5^ cells/well). THP-1 cells were differentiated into macrophages through stimulation with PMA for three days. Cells were then washed with RPMI media without antibiotics twice. For invasion, *E.coli ΔMurQ-KU* was grown and remodeled following established protocols^29,30^. 20 μL of bacteria (OD_600nm_ = 2.0) was added to each well, and the samples were incubated at 37 °C and 5% CO_2_ for 30 min. After incubation, the media was removed, and fresh media with gentamycin (1:1000) was added to kill extracellular bacteria, incubated for 30 min at 37 °C and 5% CO_2_. The media then was removed, and the cells were rinsed twice with 1xPBS at room temperature. Cells were fixed 4% paraformaldehyde in 1xPBS for 10 min at room temperature and rinsed twice with 1XPBS. Then, cells were permeabilized with 1% Triton-X in PBS for 10 min at room temperature and washed 3×5 min with 1xPBS with 0.2% tween-20 and 1.5% BSA at room temperature on a rocker. PBS (500 μL) with 0.2% tween-20 and 0.1% Triton-X was added to each well to prepare for the click reaction. To each well was sequentially added 1 mM CuSO_4_ solution, 128 μM Tris[(1-benzyl-1H-1,2,3-triazol-4-yl)methyl]amine, 1.2 mM freshly prepared (+)−sodium (L) ascorbate (Sigma-Aldrich), and 20 μM of Alk488. The click reaction was performed at room temperature for 30 min while shaking. The cells were washed 3×5min with 1xPBS with 0.2% tween-20 and 1.5% BSA at room temperature on a rocker. The cells were mounted on glass slides with 4,6-diamidino-2-phenylindole (Invitrogen) for super resolution imaging.

### Invasion assay in 3D culture for Imaging

THP-1 cells were encapsulated with an adapted version of the protocol described above. Here, cell precursor solution was prepared as before, and then 20 μL of hydrogel precursor solution was pipetted onto a 1 mL syringe mold (instead of a gasket mold) resulting in 100,000 cells per hydrogel. Hydrogels were formed by photopolymerization and placed into 24 well plates for 3D cell culture and differentiation with PMA for 3 days as before. Samples were then washed with RPMI media without antibiotics twice. For invasion, 20 μL of *E.coli ΔMurQ-KU* (OD_600nm_ = 2.0), grown and remodeled as noted above, was added to each well for 60 min, and the samples were incubated at 37 °C and 5% CO_2_. After incubation, the media was removed, and fresh media with gentamycin (1:1000) was added to kill extracellular bacteria for 60 min at 37 °C and 5% CO_2_. After 60 min, the media was removed, and the cells were rinsed twice with 1xPBS at room temperature. Cells were fixed 4% paraformaldehyde in 1xPBS for 15 min at room temperature. Fixed cells were rinsed 3×5 min with 1xPBS, and were permeabilized with 1% Triton-X in PBS for 30 min at room temperature. Cells were washed 3×5 min with 1xPBS with 0.2% tween-20 and 1.5% BSA at room temperature on a rocker. PBS (500 μL) with 0.2% tween-20 and 0.1% Triton-X was added to each well to prepare for the click reaction. To each well was sequentially added 1 mM CuSO_4_ solution, 128 μM Tris[(1-benzyl-1H-1,2,3-triazol-4-yl)methyl]amine, 1.2 mM freshly prepared (+)−sodium (L) ascorbate (Sigma-Aldrich), and 20 μM of Alk488. The click reaction was performed at room temperature for 1 hour while shaking. The cells were washed 2×30min with 1xPBS with 0.2% tween-20 and 1.5% BSA at room temperature on a rocker. The cells were washed overnight in 1xPBS with 0.2% tween-20 and 1.5% BSA at 4°C. The next day, the cells were washed 2×30min with 1xPBS with 0.2% tween-20 and 1.5% BSA at room temperature on a rocker. Cells were stained with 4,6-diamidino-2-phenylindole for 30 min and were washed 3×10min with 0.2% tween-20 and 1.5% BSA at room temperature on a rocker. Hydrogels were moved to glass chamber well slides for confocal imaging.

### SIM

Super-resolution microscopy was performed following established protocols^29,30^

### Confocal Microscopy Imaging

3D culture samples were imaged with confocal microscopy. Images were taken on Zeiss LSM800 with Plan-Apochromat 63X/1.40 Oil DIC M27 objective and frame size of 1024 × 1024 pixels. Z-stacks were 200 μm with 0.2μm slices. Excitation of 4,6-diamidino-2-phenylindole and Alk488 was achieved with 405 and 488 nm lasers, respectively. Pixel, line, and frame time were 1.03 μs, 4.95 ms, and 5.06 s, respectively. Scan direction was bidirectional, and an average of 4 scans per image was utilized. ZEN 2012 (Zeiss) was used to process the images and prepare z-stack projections.

### ELISA Preparation

Three days before stimulation with bacteria, macrophage samples were seeded and encapsulated for 2D and 3D culture, respectively. For 2D culture, cells were seeded at 1×10^6^ cells per well in a 6 well plate. For 3D culture, hydrogels were formed in gasket molds as described above, and 4 hydrogels (250,000 cells per hydrogel) were placed per well in a 6 well plate (total of 1×10^6^ cells per well). All cells were differentiated for three days with 200nM PMA. The cells were washed twice with RPMI media without antibiotics. Cells were provided RPMI media without antibiotic after the washes. The day of the stimulation with bacteria, bacterial overnights were diluted to OD = 2.0, and 20 μL of these bacteria was added to each well. To the control samples, 20μL of sterile water was added. Plates were incubated for 4 hours in an incubator (37°C, 5% CO_2_). After 4 hours, the supernatant was removed, and it was stored at −20 °C until shipment to University of Maryland Cytokine Core for analysis by ELISA.

## Results and Discussions

To fabricate a synthetic matrix with tunable properties in the range of those of the healthy gut, we utilized an 8-arm PEG functionalized with norbornene end groups (PEG-8-Nb) linked with a matrix metalloproteinase (MMP) - degradable sequence (GCRD<u>VPMS↓MRGG</u>DRCG)^31^ that is responsive to MMP-2 amongst other enzymes secreted by monocytes and macrophages.^32,33^ Additionally, these hydrogels were modified with the integrin binding peptide CGKG<u>YIGSR</u>, derived from the laminin β_1_ chain, to promote cell adhesion inspired by the laminin-rich ECM of the basement membrane of the gut.^34,35^ The hydrogels were formed rapidly through light-triggered thiol-ene click chemistry via a step growth mechanism, as the norbornene groups on the PEG are coupled to the thiols presented by the cysteines in the difunctional MMP-degradable linker peptide and monofunctional integrin binding peptide^36^ (Figure 2A and B). The reaction was initiated using the photoinitiator lithium phenyl-2,4,6-trimethylbenzoylphosphinate (LAP) and low, cytocompatible doses of light (10 mW/cm^2^ at 365 nm for 2 min). These synthetic bioinspired hydrogels were designed with a Young’s modulus (E ~ 2.6 ± 0.8 kPa) to mimic the ‘stiffness’ of the healthy gut^37^, as confirmed by shear rheometry (Figure 2C).

**Figure 2:**
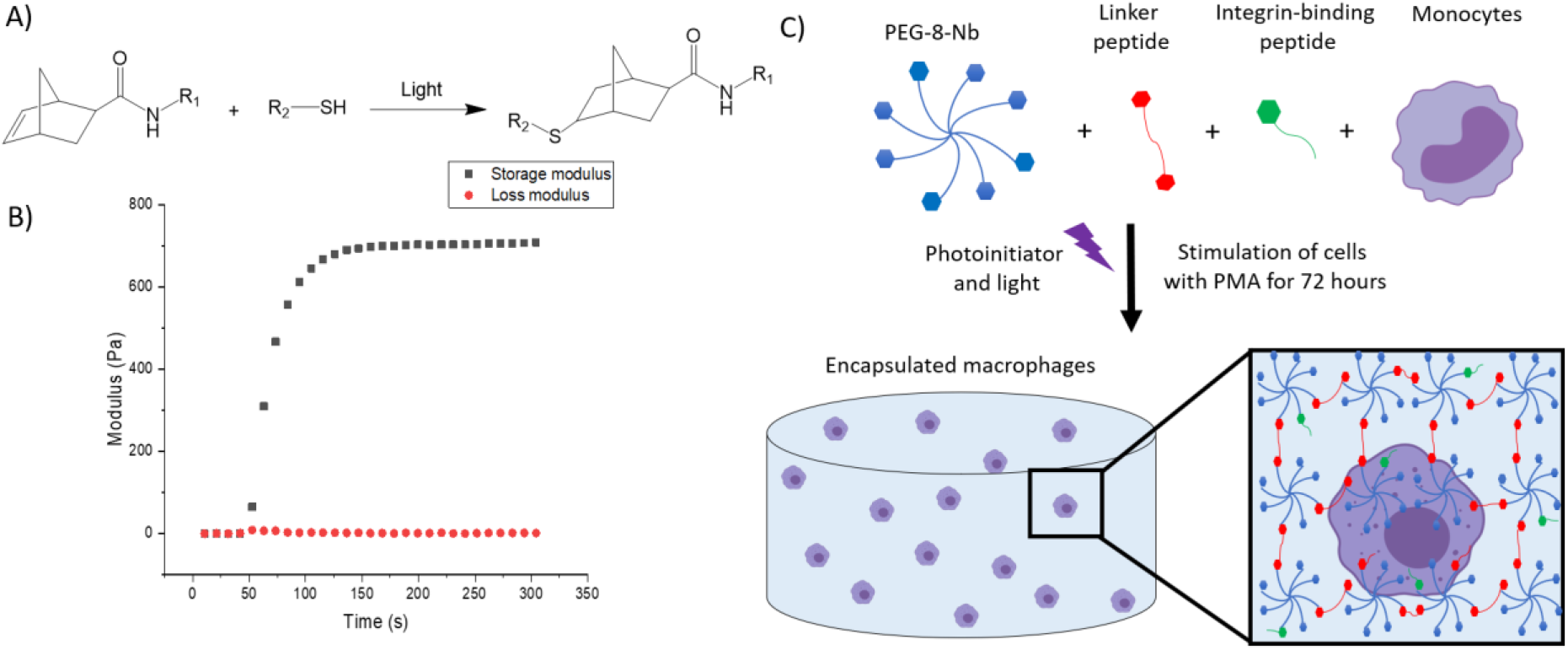
Overview of hydrogel design. **A)** Hydrogels were formed using thiol-ene click chemistry with norbornenes on the PEG and thiols on the difunctional linker and monofunctional integrin binding peptides. **B)** Hydrogels are rapidly formed upon the application of light. The increase in storage modulus, monitored with *in situ* rheometry, indicates crosslinking events, and complete hydrogel formation was observed within 2 minutes. This final modulus, converted to Young’s modulus by rubber elasticity theory for ease of comparison to native tissues, is relevant for mimicking the modulus or ‘stiffness’ of the healthy gut. **C)** The hydrogel network is composed of 8-arm PEG-norbornene (blue), thiol-containing linker peptide (red), and thiol-containing integrin binding peptide (green). Components were photo-polymerized with a low, cytocompatible dose of light (10 mW/cm^2^ at 365 nm for 2 min) to form a hydrogel, and the monocyte cells in the hydrogel subsequently were differentiated into macrophages through stimulation with PMA for 72 hours. Schematic not to scale.

This type of PEG-peptide hydrogel has been used previously to culture a wide variety of cell types, including human mesenchymal stem cells, induced pluripotent stem cells, and breast cancer cells.^13,25,38,39^ However, as the mechanism of hydrogel polymerization involves free-radicals, the viability of sensitive cell types can be impacted during cell encapsulation and hydrogel formation.^31^ Accordingly, we examined cell viability upon encapsulation and during 3D cell culture within these materials. Here, THP-1 cells were selected as a human monocytic cell line that are commonly are differentiated into macrophage cells, as human primary tissue macrophages cannot be readily expanded *ex vivo.*^40^ We first differentiated monocytes into macrophages on TCPS and subsequently encapsulated these cells. We observed low viability in the hydrogels (SI Figure S1). Therefore, we examined encapsulating monocytes and then differentiating them into macrophages within the matrix, which also mimics aspects of the natural process of monocyte arrival and differentiation into macrophages within the native gut. Importantly, monocytes were successfully encapsulated within the hydrogels and exhibited high viability throughout 3D culture (Figure 3A, SI Figure S2).

**Figure 3:**
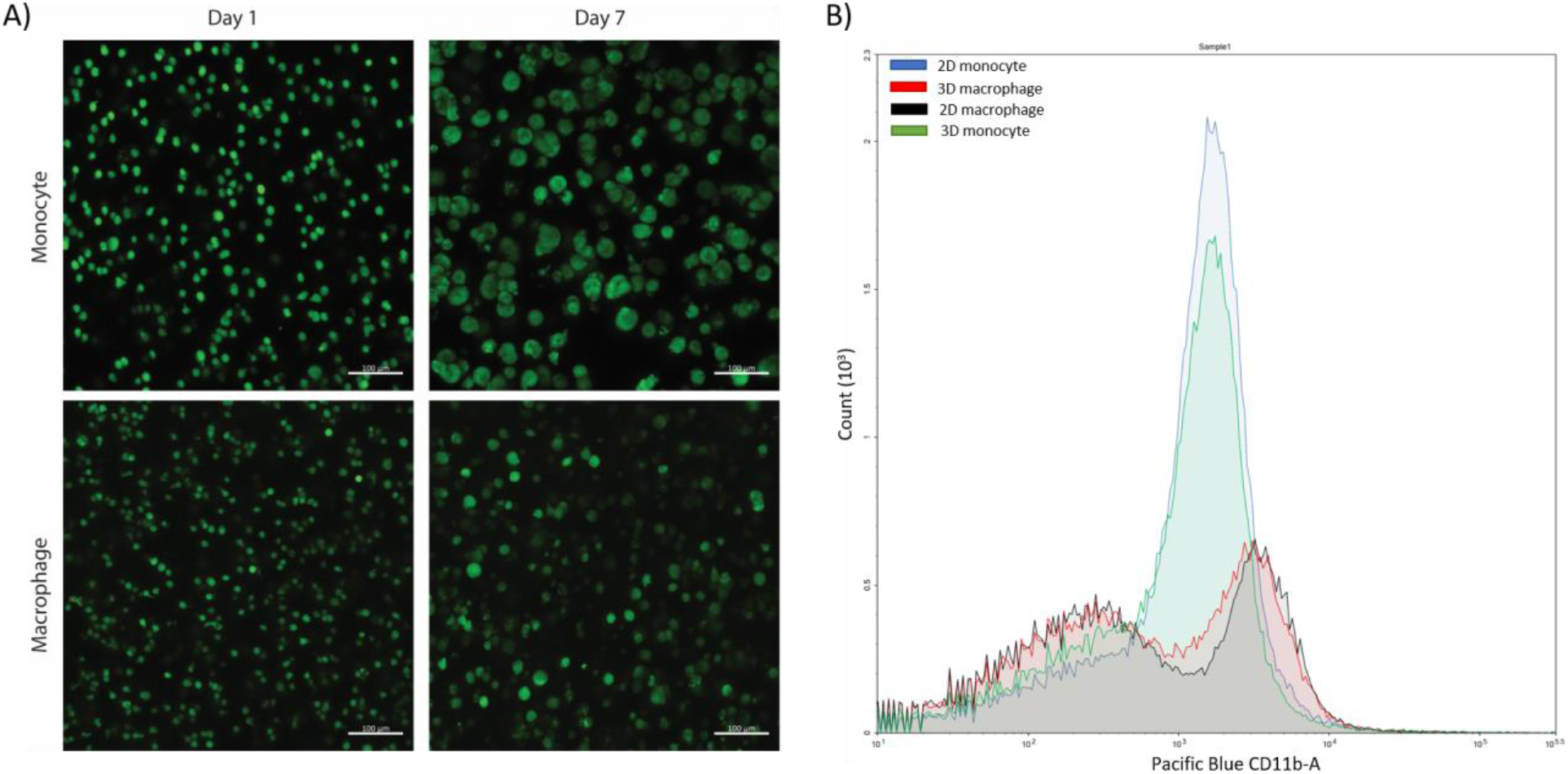
Confirmation of cell viability and cell differentiation in 3D culture. **A)** Live/dead cytotoxicity results of THP-1 cells (monocytes) and THP-1 cells differentiated with PMA (macrophages) in PEG-peptide hydrogels. Images are representative of a minimum of three fields per view per hydrogel of three biological replicates. **B)** Flow cytometry data of macrophages and monocytes in 2D or 3D culture stained for extracellular macrophage marker CD11b. 2D (blue) and 3D monocyte (green) profiles match each other, and 2D (red) and 3D (black) macrophage profiles match each other, showing differentiation is achieved in the 3D hydrogel. Plot is representative of 3 biological replicates and two technical replicates.

After confirmation of successful culture of THP-1 monocyte cells, monocytes were differentiated into macrophage cells through stimulation with phorbol 12-myristate 13-acetate (PMA) over three days based on literature precedent in two dimensions.^40–42^ Differentiation was confirmed through hydrogel digestion and subsequent staining and flow cytometry of recovered cells for macrophage extracellular and intracellular markers, CD11b and CD68, respectively (Figure 3B, SI Figure S3). Further, these macrophages also exhibited high viability in the hydrogels (Figure 3A, SI Figure S2).

With successful 3D culture of macrophages, we focused on modeling invasion with fluorescently labeled bacterial cell walls. Work has been done previously to label various aspects of PG including small fluorophores coupled to D-amino acids, NAG and NAM sugar probes, and NIR fluorogenic probes.^14^ We chose to utilize the NAM probe previously developed in our lab because synthetic fragments MDP and MTP both contain the NAM sugar, leading us to hypothesize that NAM plays an important structural role in fragment identification and subsequent infection clearing by macrophages. Additionally, the NAM probe has a bioorthogonal azide at the 2-acetyl position, providing versatility in what is ‘clicked’ onto the handle for various assays.^29,30^ Therefore, utilizing these NAM probes can allow us to visualize the bacterial core and have potential downstream applications as an affinity handle for pulling out naturally produced PG fragments from breakdown in macrophages. Another advantage of the NAM probe is that under a lethal dose of Fosfomycin, MurA, the first enzyme in the PG biosynthetic pathway, can be selectively inhibited, which prevents natural PG biosynthesis from occurring, and cells only survive if they uptake the 2-azido NAM sugar provided.^43^ Instrumental work by Mayer and coworkers showed that in certain species of bacteria, such as the *Pseudomonas* family, there is recycling machinery involving the enzymes anomeric NAM/NAG kinase (AmgK) and α-1-phosphate uridylyl transferase (MurU) that under a lethal dose of Fosfomycin can recycle NAM into uridine diphosphate (UDP)-NAM, which is then incorporated into mature PG.^44^ A previously engineered strain of *E. coli* (*E. coli ΔMurQ KU*) that expresses recycling enzymes AmgK and MurU was used to specifically label the NAM residue of PG.^29^ We wanted to assess if the macrophages in 3D culture within hydrogels were amenable with this system.

Through remodeling with the 2-azido *N*-acetyl muramic acid sugar (2Az-NAM), we observed fluorescent labeling specifically to the cell wall of the *E. coli* PG following the copper catalyzed azide-alkyne cycloaddition (CuAAC) click reaction to install an alkyne fluorophore (Alk488) (Figure 4). With the ability to fluorescently label *E. coli*, macrophages were invaded with bacteria on 2D culture on TCPS and into 3D cultures within synthetic hydrogels. After invasion for 1 hour, we observed engulfment of the bacteria by the macrophages in both 2D and 3D culture (Figure 4, SI Figure S4), indicating that this system will have utility in identifying naturally released bacterial peptidoglycan fragments.

**Figure 4:**
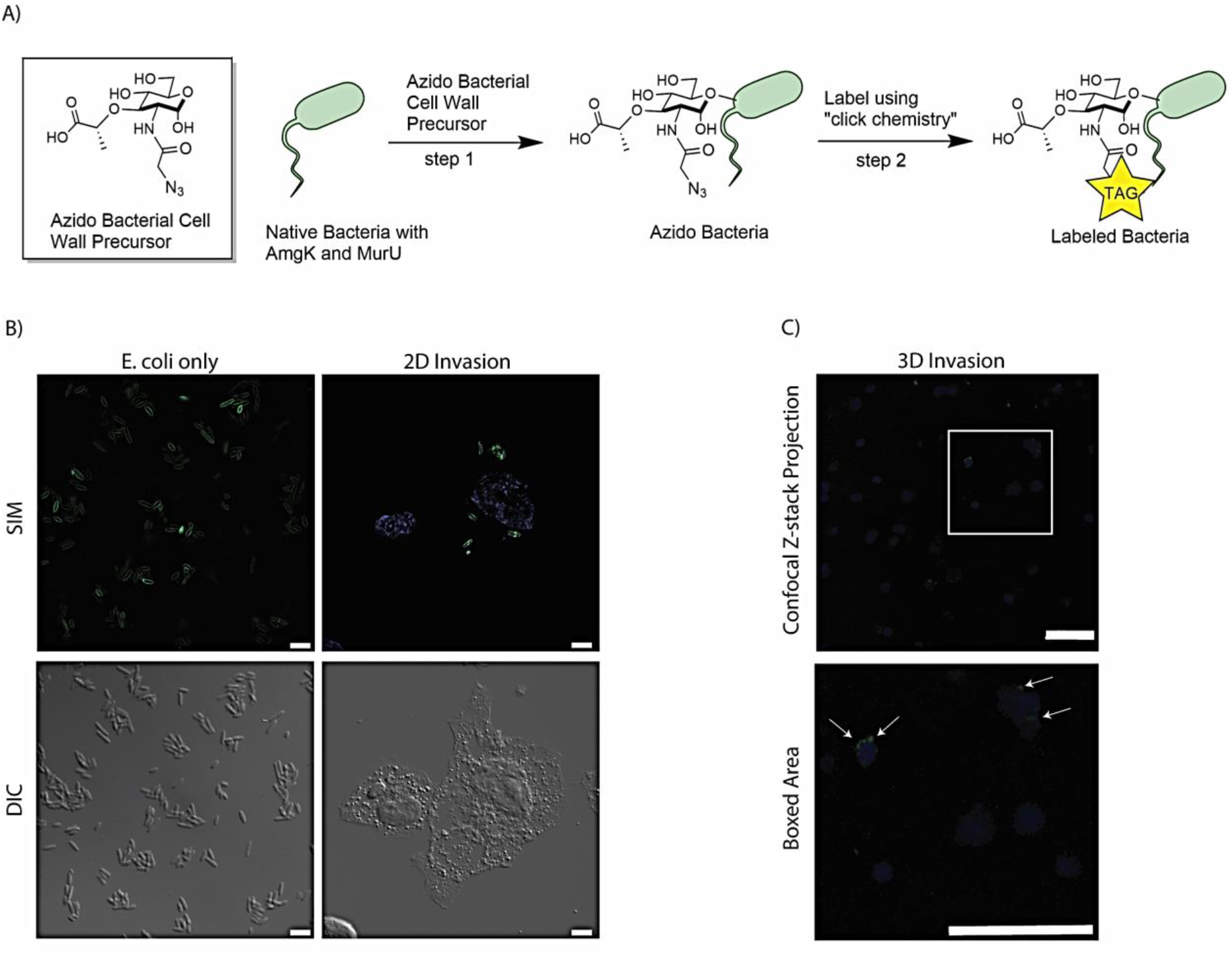
THP-1 macrophage cells invaded by remodeled *E. coli ΔMurQ KU* in 2D and 3D culture. **A)** *E. coli ΔMurQ KU* cells are remodeled by incorporating the azido bacterial cell wall precursor in cell wall with recycling enzymes AmgK and MurU. Bacteria are then fluorescently labeled using CuAAC. **B)** Differential interference contrast (DIC) and individual 2D images from super-resolution structured illumination microscopy (SIM) of *E. coli ΔMurQ KU* cells remodeled with 2Az-NAM. *E. coli ΔMurQ KU* cells were invaded into THP-1 macrophage cells on glass coverslips. All cells were fixed and Alk488 was clicked on to the remodeled bacteria (green). Nuclei were labeled with 4,6-diamidino-2-phenylindole (blue) (scale bars 5 μM). **C)** Confocal microscopy z-stack projections of *E. coli ΔMurQ KU* cells invaded into THP-1 macrophages cells encapsulated in hydrogel-based synthetic matrix for 3D culture. All cells were fixed and Alk488 was clicked on to the remodeled bacteria (green). Nuclei were labeled with 4,6-diamidino-2-phenylindole (blue) (scale bars 50 μM). Images are representative of a minimum of three fields viewed per replicate with at least two technical replicates, and invasion experiments were conducted in at least three biological replicates.

Following successful invasion of macrophages in both 2D and 3D culture, we aimed to probe potential differences in cytokine expression as a measure of macrophage activation upon invasion, which may be influenced by the dimensionality and complexity of the culture system. Here, we utilized the *E. coli ΔMurQ KU* strain as noted before and another Gram-negative organism, *Pseudomonas aeruginosa*. *P. aeruginosa* is an opportunistic, pathogenic species of bacteria that is known to break through mucosal barriers particularly in hospital infections and is extremely antibiotic resistant.^45^ To assess macrophage response, we examined secretion of cytokines associated with different aspects of macrophage activation. Specifically, an enzyme-linked immunosorbent assay (ELISA) was implemented to observe cytokine output of tumor necrosis factor alpha (TNF-α) and interleukin 6 (IL-6) of cells treated with *E. coli*, *P. aeruginosa*, or no bacteria. TNF-α is a cytokine involved in systemic inflammation and regulates immune cells, induces fever, inhibits viral replication, and induces apoptosis. Dysregulation of TNF-α is implicated in various diseases including IBD. IL-6 is a cytokine involved in stem cell differentiation, antibody synthesis by B cells, and T cell cytotoxicity.^46^ IL-6 is produced in response to bacterial and viral infections, and work has shown that dysregulation of IL-6 can be involved in autoimmune disease pathogenesis.^46^

When macrophages were invaded with either *E. coli* or *P. aeruginosa*, an increase in TNF-α expression was observed, which accurately represents known responses THP-1 macrophages have to lipopolysaccharide (LPS) located on Gram-negative bacteria^42,47^ (Figure 5). Further, THP-1 macrophage cells from 2D plates treated with *E. coli* showed statistically significant increased expression of IL-6 as compared to untreated cells and cells treated with *P. aeruginosa*. Notably, THP-1 macrophage cells treated with *E. coli* in 3D culture did not show statistically different expression of IL-6 compared to the control and *P. aeruginosa* samples (Figure 5). These data suggest that the 3D invasion model produced a more biologically relevant response of IL-6 to the various bacterial species, as THP-1 macrophages treated with LPS are known to have little IL-6 response.^47^

**Figure 5:**
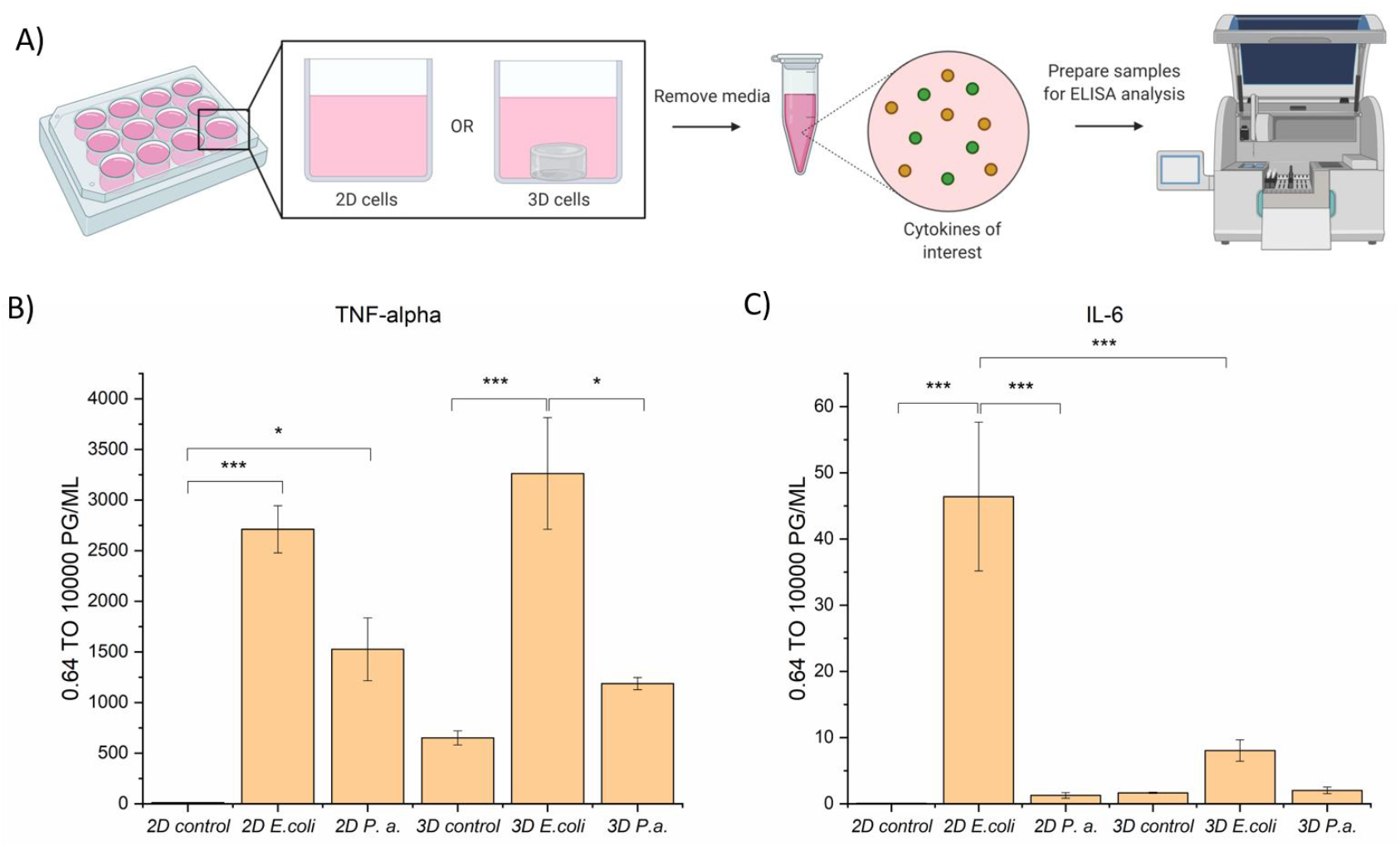
ELISA analysis of TNF-α and IL-6 cytokine production by THP-1 macrophage cells in 2D and 3D culture upon invasion with *E. coli* or *P. aeruginosa*. **A)** Macrophage cells in 2D or 3D culture were stimulated with bacteria or water for 4 hours. Cell culture supernatant was harvested and subjected to ELISA analysis for TNF-alpha and IL-6 cytokines. **B)** ELISA analysis of TNF-alpha shows cells stimulated with *E. coli* or *P. aeruginosa* exhibited increased secretion of TNF-alpha as compared to the control. **C)** ELISA analysis of IL-6 shows cells stimulated with *E. coli* in 2D exhibited increased secretion of IL-6 as compared to in 3D culture and the controls. Macrophages in 2D and 3D culture when stimulated with *P. aeruginosa* did not exhibit increased secretion of IL-6 as compared to the control. All samples were analyzed in biological triplicate and technical duplicate and statistically analyzed by ANOVA with Tukey’s test (*p < 0.05, **p<0.01, ***p<0.001).

## Conclusion

Overall, we have established a well-defined hydrogel-based system for the 3D culture of monocytes and macrophages in an environment that mimics aspects of the basement membrane of the gut, and a multi-dimensional invasion assay for probing macrophage response to invading bacterial species. The system is reproducible, robust, and based on the underlying design of the synthetic matrix, provides future opportunities for not only probing the impacts of host-bacterial cell interactions, but also cell-matrix interactions through manipulation of extracellular biochemical and mechanical cues. The system is amenable to bioorthogonal chemistry on bacterial peptidoglycan within the hydrogel and opens the door for further applications in chemical biology involving labeling and study of the bacterial cell wall. This model system can be further tailored for studying cell-cell and cell-matrix interactions in disease progression and for asking new questions beyond traditional 2D culturing methods that prevail in the field. The use of the NAM probes with the azide bioorthogonal handle embedded in the bacterial peptidoglycan will provide opportunities to enrich for biologically produced bacterial cell wall fragments from these invasion systems for further analysis. These studies provide new insights into the interactions between bacteria and immune cells, and this new model system provides a platform to examine how our immune system recognizes bacteria in a variety of physiologically relevant settings.

## Acknowledgements

We thank Dr. Jeffery Caplan, the Director of Bioimaging, and the Delaware Biotechnology Institute for assistance with super-resolution microscopy. We thank Lisa Hester, and the University of Maryland Cytokine Core Laboratory, for assistance with ELISA assays. We thank Dr. Catherine Fromen and graduate student Bader Jarai for access and training on flow cytometry. K.A.W. would like to thank Dr. Lina Pradhan for initial training and mentorship for hydrogel chemistry and Dr. Kristen DeMeester for initial training and mentorship for bacteria labeling and mammalian cell culture. We would like to thank Elizabeth D’Ambrosio and Samantha Cassel for critical reading of this manuscript. We are thankful for support from the Delaware COBRE program, supported by a grant from the National Institute of General Medical Sciences (NIGMS 1 P30 GM110758 and 1 P20 GM104316-01A1). This research was supported by grants for related work from the NIH U01 Common Fund program with grant number U01CA221230-01 (C.L.G.) and the NIH Director’s New Innovator Award with grant number DP2HL152424 (A.M.K). K.A.W. would also like to thank the NIH for support through Chemistry-Biology Interface (CBI) training grant, 5T32GM008550. TOC and Figure 5 were created using BioRender.com.

