## Supporting Information for "Multiscale Invasion Assay for Probing Macrophage Response to Bacteria"

### Table of Contents

I. **General Materials and Methods**……………………………………………………….…3
II. **Synthetic Methods**…………………………………………………………………....3
III. **Mammalian Cell Culture**…………………………………………………………….5
IV. **Live/Dead Viability Assay on Encapsulated Cells**…………………………………..…5
V. **Bacterial Labeling Protocol**…………………………………………………………..…5
VI. **Live/Dead of Macrophages Differentiated on Plate**………………………………..7
VII. **Fraction Viable of Monocytes and Macrophages Differentiated in the Hydrogel**…………………………………………………………………….…...8
VIII. **Flow Cytometry Data for CD68 Macrophage Marker** ………………………………9
IX. **Invasion of *E. coli* into Macrophages in 3D culture**…………………………………..10
X. **MS Spectra of Peptides**…………………………………………………….……...11
XI. **NMR Spectra of PEG-8-Nb** ……………………………………………………....13
XII. **References**……………………………………………………………………….…14

#### I. General Materials and Methods

**Materials.**

All chemicals were purchased from Millipore-Sigma and ThermoFisher Scientific and used without further purification unless otherwise noted. Antibiotics were purchased from Gold Biotechnology. Deuterated NMR solvents were purchased from Cambridge Isotope Laboratories. *E. coli ΔMurQ-KU* strain was developed in the Grimes Lab[1]. THP-1 cells were purchased from American Type Culture Collection (ATCC). Cell culture supplies were purchased from Cell Treat.

**Instrumentation.**

NMR spectra were recorded on AV III 600 MHz spectrometers at the University of Delaware NMR Facility. Mass Spectrometry (LRMS, ESI) were obtained using an ACQUITY UPLC H-Class/SQD2 at the University of Delaware Mass Spectrometry Facility. Peptides were synthesized using an automated peptide synthesizer (PS3, Protein Technologies) and were purified on reverse-phase high performance liquid chromatography (HPLC, XBridge BEH C18 OBD 5 μm column; Waters, Milford, MA). Rheometry was performed on AR-G2 rheometer with UV-visible light attachment (TA instruments) in tandem with an Omincure Series 2000 light source (Excilitas) with a 365 nm bandpass filter and light guide (Exfo). Confocal microcopy images were taken an Images Zeiss LSM800 instrument with Plan-Apochromat 63X/1.40 Oil DIC M27 objective. Super-resolution microscopy images were taken on Zeiss Elyra PS. 1 microscope with Plan-Apochromat 63x/1.4 oil differential interference contrast (DIC) M27 objective.

#### II. Synthetic Methods

**Synthesis and characterization of norbornene-functionalized PEG.**

Multi-arm (8) PEG ($M_n$ ~ 40,000 g/mol) was functionalized with norbornene end groups according to established protocols[2,3]. Briefly, in a 250 mL round bottom flask, PEG-8-$NH_2$·HCl (5 g, Jenkem) was dissolved in anhydrous *N*,*N*-dimethylformamide (DMF, ThermoFisher) and stirred at room temperature. In a second 250 mL round bottom flask, 5-norbornen-2-carboxylic acid (Nb-COOH) (17.6 molar equivalent; 2.2 excess relative to amine groups on the PEG), 4-methylmorpholine (4-MMP) (36 molar equivalent), and HATU (16 molar equivalent; 2 excess) were dissolved in 10 mL of DMF stirring at room temperature. Once the individual flask components were dissolved, they were combined into one flask and stirred at room temperature overnight. The solution was then precipitated twice in cold diethyl ether (500 mL, 14x excess diethyl ether relative to DMF), and the resulting suspension was filtered with a Buchner funnel with filter paper to recover the precipitated polymer product. The solid PEG product was dried in the vacuum oven overnight. The PEG was purified by dialysis (MWCO 1000 g/mol) in mq$H_2$0 for 48 hours. Product purity was confirmed by $^1$H NMR in DMSO-$d_6$. Norbornene functionality was determined to be approximately 75% on average (Section XI below), and PEG-8-Nb was stored at -20°C following lyophilization.

**Synthesis and characterization of peptides.**

All peptides were synthesized using solid phase peptide synthesis based on established protocols[2]. The difunctional linker peptide (GCRDVPMSMRGGDRCG) and the monofunctional pendant peptide (CGKGYIGSR) were synthesized using standard FMOC-chemistry on an automated peptide synthesizer (PS3 Peptide Synthesizer; Protein Technologies, Inc., Tucson, AZ). The peptides were built on Rink Amide MBHA resin. All amino acids were double coupled. The peptides were cleaved from the resin for 2-3 hours in 95% trifluoracetic acid, 2.5% water, and 2.5% triisopropylsilane supplemented with 50 mg/ml dithiothreitol. Following cleavage, all peptides were precipitated in cold diethyl either at 4°C and let airdry overnight. The peptides were purified by reverse-phase high performance liquid chromatography (HPLC; XBridge BEH C18 OBD 5 μm column; Waters, Milford, MA) with a linear 95%/5 % to 5% /95% linear water-acetonitrile (ACN) gradient over 15-30 min. Purified peptides were subsequently lyophilized. Their molecular weights were verified by mass spectrometry (Section X below), and the thiol concentration of each peptide was determined using Ellman's assay. Purified peptides were dissolved in phosphate buffered saline (PBS) and stored at -80°C.

**Synthesis and rheological characterization of hydrogels.**
Monomer stocks were prepared by dissolving each component in phosphate buffered saline (PBS): PEG-8-Nb (40 mM Nb functionality); lithium phenyl-2,4,6-trimethylbenxoylphospohinate (LAP) (30 mM) sterile filtered with 0.45 μM filter; and each peptide (~200 mM). PEG-8-Nb and LAP stocks were stored at -20°C, and peptide stocks were stored at -80°C.

A bulk hydrogel precursor solution was prepared using 7mM PEG-8-Nb, 5mM peptide crosslinker, 2mM pendant peptide, and 2mM LAP in PBS. Rheology measurements were conducted as previously reported[2] on AR-G2 rheometer with UV-visible light attachment (TA instruments) in tandem with an Omincure Series 2000 light source (Excelitas) with a 365 nm bandpass filter and light guide (Exfo). Briefly, 10 µL of hydrogel precursor solution was pipetted on to the quartz plate of the UV-vis light attachment on the rheometer with a 8mm flat plate geometry installed, and the gap was set to 150 µm. Hydrogel crosslinking and gelation were monitored by measuring storage (G') and loss (G") moduli at 0.5% applied strain and 2 rad/s frequency upon irradiation (10 mW/cm$^2$ at 365 nm). The gelation time was determined to be less than 2 minutes based on the change in G' being within 5% between two consecutive points[4]. Frequency sweeps at 1% strain were performed after the irradiation was complete to measure the final moduli of hydrogels formed *in situ*. All of the rheometric measurements were performed within the linear viscoelastic regime. Final equilibrium swollen moduli at physiological temperature were calculated as previously reported[3,5] using the *in situ* measured modulus and the following equations:

$$G_{final} = G_0\left(\frac{T_{final}}{T_0}\right)\left(\frac{Q_{final}}{Q_0}\right)^{-\frac{1}{3}}$$
$$Q_{final} = (1-2\chi)N^{0.57}\phi^{-0.38}$$

where $G_0$ is the *in situ* measured shear modulus; $T_{final}$ is 310 K; $T_0$ is 298 K; $Q_0$ is the initial volumetric swelling ratio of the hydrogels; χ is 0.426, the PEG–water interaction parameter; N is 304, the number of PEG repeats between crosslinks; and ϕ is the initial volume fraction of polymer. Lastly, the Young's modulus (E) was estimated by rubber elasticity theory:

$$E = 2G_{final}(1+\nu)$$

where ν is Poisson's ratio and is taken to be 0.5 for these incompressible, elastic PEG hydrogels.

#### III. Mammalian Cell Culture

THP-1 cells were purchased from the American Type Culture Collection (ATCC) and cultured under sterile conditions at 37 °C with 5% $CO_2$. Cells were grown in RPMI media, 10% FBS (Atlantic Biologicals), 2 mM L-glutamine, and 2 mM penicillin-streptomycin.

#### IV. Live/Dead Viability Assay on Encapsulated Cells

THP-1 monocyte and macrophage cell viability following encapsulation in the hydrogels was assessed on days 1, 3, and 7 using a LIVE/DEAD® Viability/Cytotoxicity Kit (ThermoFisher Scientific). THP-1 monocytes were encapsulated as a single cell suspension at a density of $5x10^6$ cells per mL in 20 µL of hydrogel precursor solution (250,000 cells/hydrogel). Precursor solution was prepared using 7mM PEG-8-Nb, 5mM linker peptide, 2mM pendant peptide, 2mM LAP, and THP-1 cell suspension in PBS. Hydrogels were formed in 10 mm x 0.5 mm sterile gasket molds (43 µL precursor solution/mold) upon irradiation with a cytocompatible dose of long wavelength UV light (10 mW/$cm^2$, 365 nm, 2 min; Omnicure 2000 with light guide and collimating lens). Two replicates were formed at a time and then placed in a 24 well plate with 500 µL of RPMI media. 200nM TPA was added to the media in each well for differentiation into macrophages. Cell-hydrogel constructs were incubated under sterile conditions at 37°C with 5% CO2. Hydrogels (n = 3) were removed from incubator on days 1, 3, and 7, and washed 2x5 min with 500 µL of PBS followed by a 20-minute incubation (37 °C at 5% CO2) with 500 µL of PBS containing calcein AM (2 µM) and ethidium homodimer-1 (4 µM). After staining, hydrogels were again washed (2x5 min with 500 µL of PBS) before imaging. Hydrogels were transferred to a chamber slide (Nunc Lab-Tek™ II Chamber Slide, Glass, 1 well) and imaged with confocal microscopy (Zeiss LSM 800, 10x objective at a zoom of 0.6x and frame size of 1024 × 1024 for each image, 200 µm z-stack, 3 images per hydrogel sample). Orthogonal projections were made of each z-stack, and live (green) and dead (red) cells were counted using ImageJ.

#### V. Bacterial Labeling and Imaging

**Bacterial Remodeling and Labeling.**
Bacterial remodeling and labeling was achieved based on established protocols for *E. coli*[1,6]. Briefly, overnight pre-cultured *E. coli ΔMurQ-KU* cells[1] were inoculated into fresh LB medium and were incubated until the $OD_{600nm}$ was about 0.600. 1.2 mL of cells were collected by centrifugation at 8,000 rpm (6,000 g) for 5 min. *E. coli ΔMurQ-KU* cells were resuspended in 190 µL LB medium. 6mM NAM sugar 1 or 2 (NAM, AzNAM, 1 mM isopropyl-1-thio-β-D-galactoside) and 200 µg/ml fosfomycin were added into all cell samples. Cells were incubated while shaking at 37 °C for 60 min. Cells were then collected (10,000 rpm, 2 min) and washed with 2x600 µl 1xPBS. Cells were fixed in 4% paraformaldehyde in PBS for 20 min. Cells were washed with 2x600 µl 1xPBS. Cells were resuspended in 190 µl PBS to prepare for the click reaction. To the bioorthogonally tagged bacterial cells was sequentially added 1 mM $CuSO_4$ solution, 128 µM Tris[(1-benzyl-1H-1,2,3-triazol-4-yl)methyl]amine, 1.2 mM freshly prepared

(+)−sodium (L) ascorbate (Sigma-Aldrich) and 20 μM of Alk488. Cells were incubated at room temperature for 30 min. Cells were washed 2x600 μl 1xPBS, 1x200 μl 1xPBS, 1x200 μl 1xPBS for 45 min in the dark, and 1x200 μl 1xPBS. The cells were resuspended in 100 μl 1xPBS and prepared for imaging.

**Structured Illumination Microscopy (SIM).**
Bacterial labeling was imaged on a Zeiss Elyra PS. 1 microscope with Plan-Apochromat 63x/1.4 oil differential interference contrast (DIC) M27 objective. Excitation of Alk488 was achieved with 488 nm laser excitation, and the camera exposure time was set to 100.0 ms. The raw data contained 5 rotations with 0.110 μm z-stack interval. Images were processed in Carl Zeiss ZEN 2012 to construct SIM images. Processing and filtering settings were kept constant and image intensity was preserved using the raw image scale option. Two-dimensional (2D) SIM images and 2D maximum intensity projection images were generated with Zen 2012.

#### VI. Live/Dead of Macrophages Differentiated on Plate and Subsequently Encapsulated

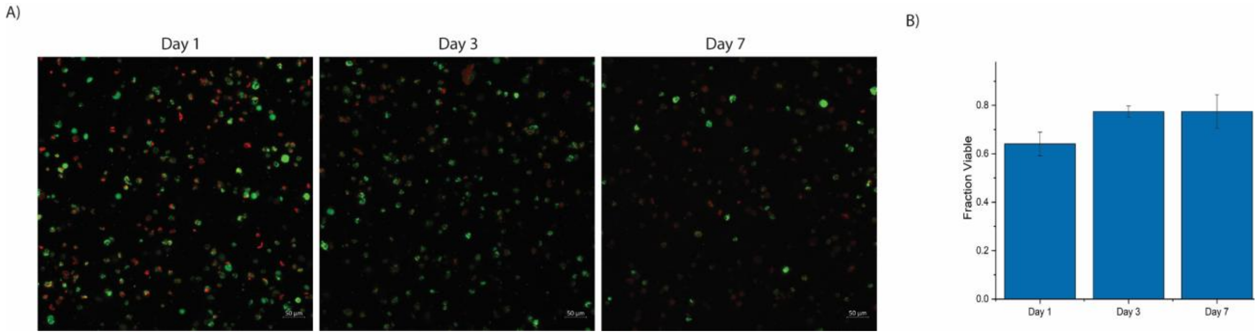


**SI Figure 1:** A) Representative images of live/dead cytotoxicity assay results of THP-1 cells differentiated with PMA into macrophages on tissue culture polystyrene and subsequently encapsulated within hydrogels. B) Quantification of fraction viable in A showed viability less than 80%, which was viability observed when THP-1 monocytes were encapsulated in the hydrogels and subsequently differentiated (see SI Figure 2).

#### VII. Fraction Viable of Monocytes and Macrophages Differentiated in the Hydrogel

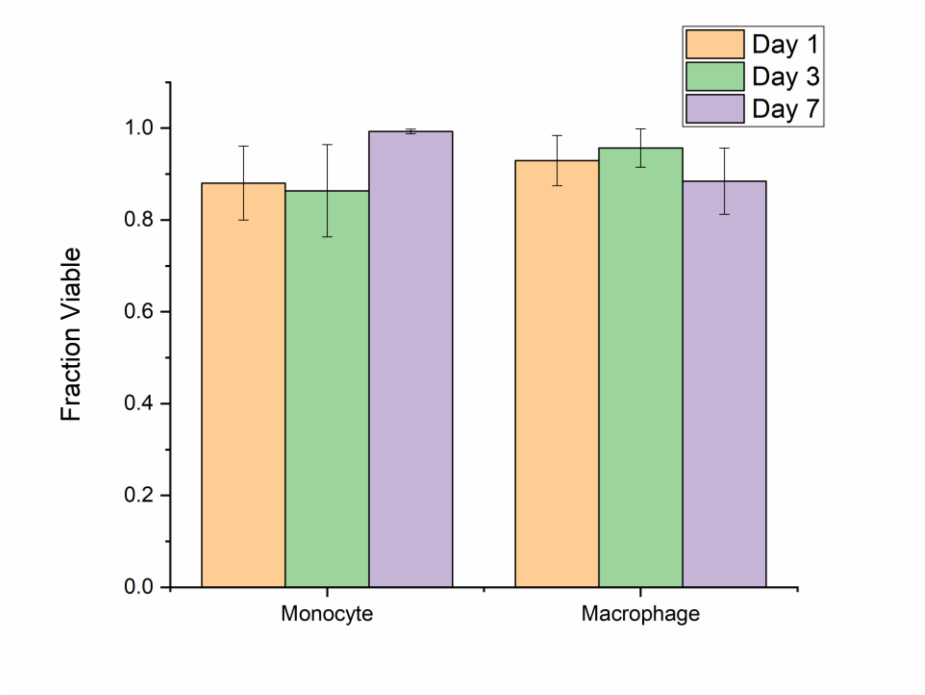


**SI Figure 2:** Quantification of fraction viable THP-1 monocyte cells in 3D culture and THP-1 macrophage cells, which were encapsulated as monocytes and subsequently differentiated with TPA during 3D culture, showed high viability for both monocytes and macrophages in 3D culture (above ~ 85% and 90%, respectively) within hydrogels.

#### VIII. Flow Cytometry Data for CD68 Macrophage Marker

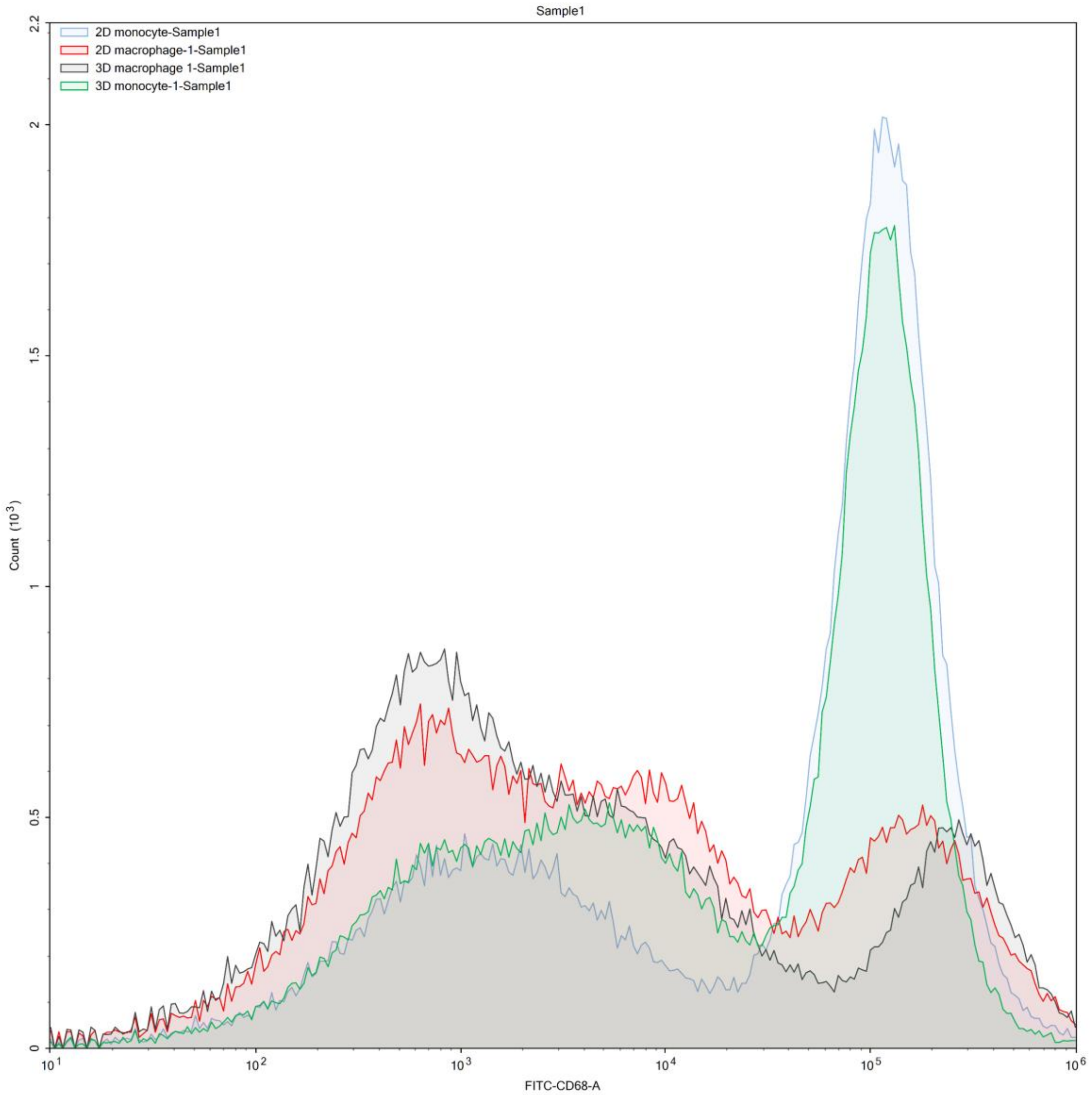


**SI Figure 3:** Flow cytometry data of macrophages and monocytes in 2D and 3D culture stained for extracellular macrophage marker CD68. 2D (blue) and 3D culture monocyte (green) profiles match each other, and 2D (red) and 3D (black) culture macrophage profiles match each other, showing differentiation was achieved in 3D culture within hydrogels. This data supports the data shown in the main text for macrophage marker CD11b (Figure 3).

#### IX. Invasion of *E. coli* into Macrophages in 3D culture

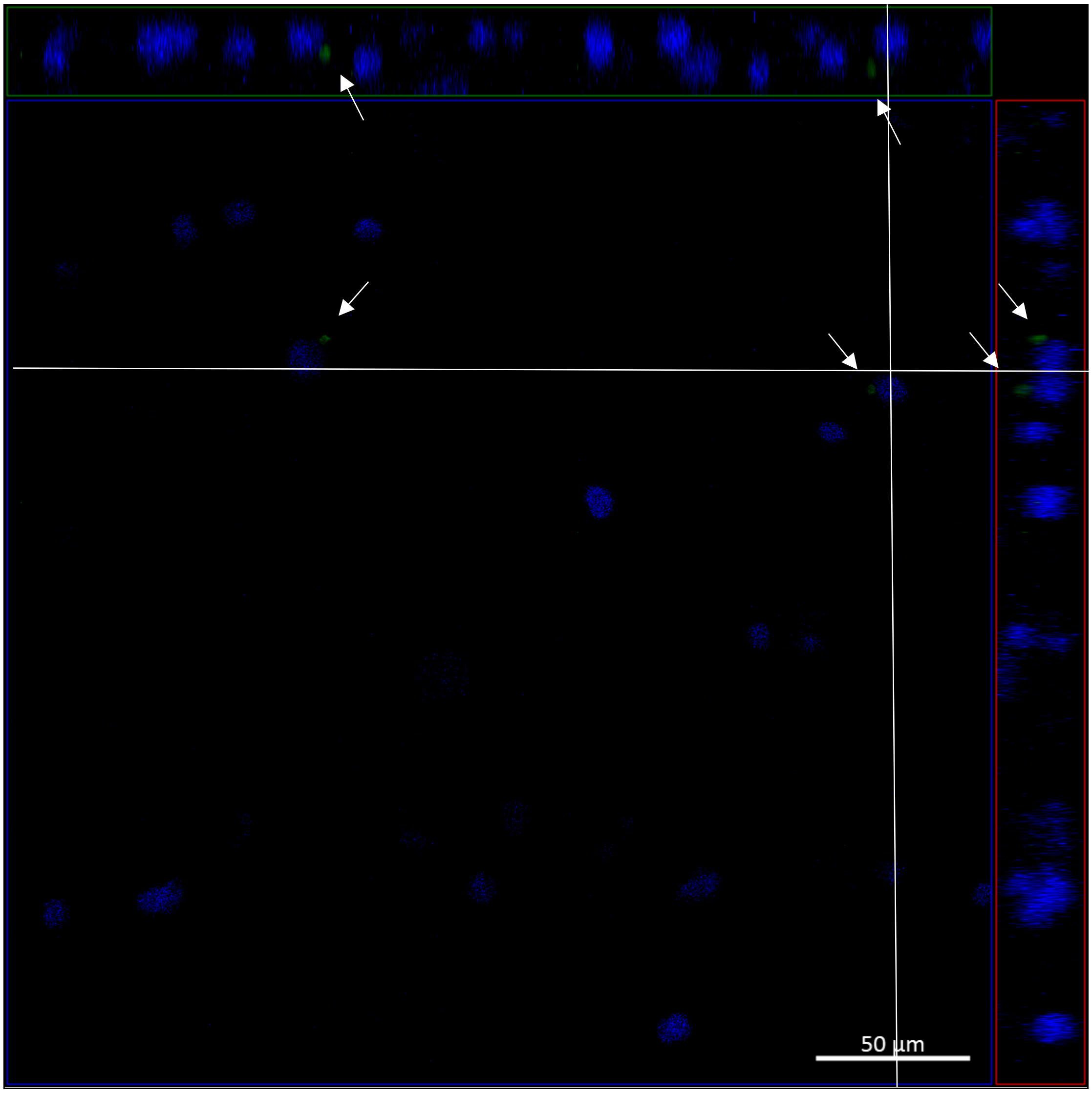


**SI Figure 4:** Representative maximum intensity projection for visualizing the invasion of E. coli into macrophages in 3D culture. The blue square is the XY orthogonal projection; the green rectangle is an XZ projection; and the red rectangle is the YZ projection. Macrophages are stained with DAPI (blue) for DNA to label nuclei, and bacteria are shown in green (labeled with AF488). Image is representative of at least 3 fields of view for each of the three biological replicates.

#### X. MS of Peptides

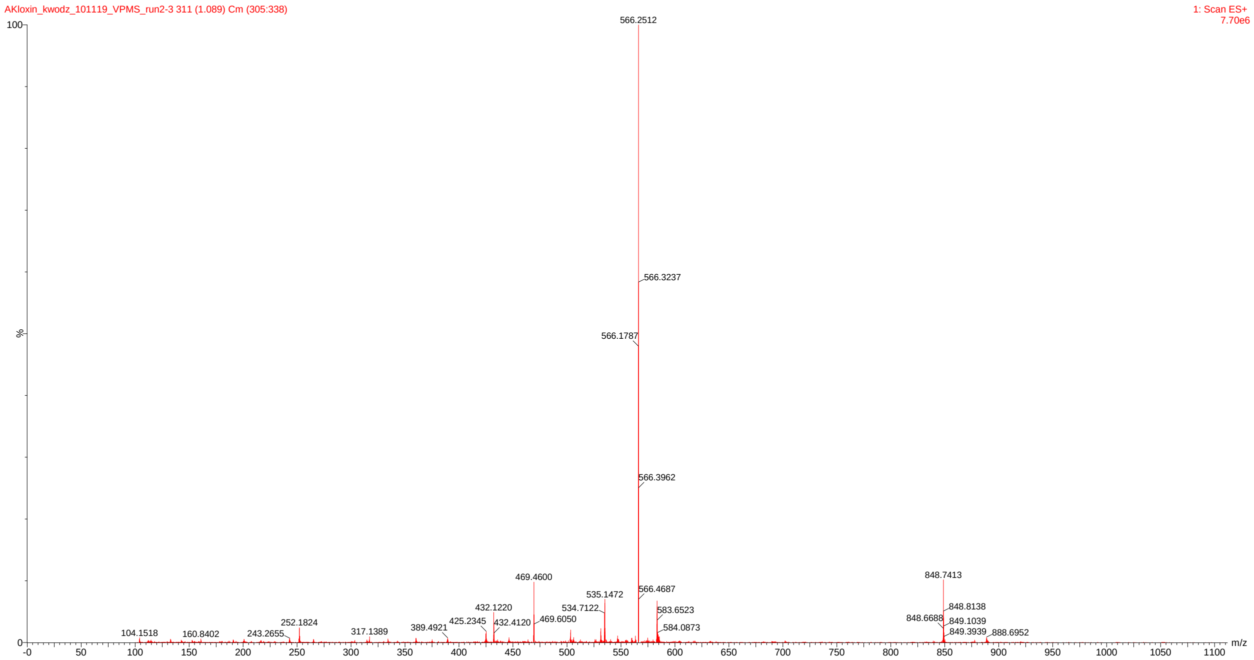


**SI Figure 5:** Mass spectrometry of difunctional linker peptide. Successful synthesis of crosslinking peptide, CGRDVPMSMRGGDRCG-amide, was confirmed by SQD2 mass spectrometry. Expected molecular weight of 1696 g/mol. $[M + 2H]^+$ = 848 g/mol $[M + 3H]^+$ = 566 g/mol.

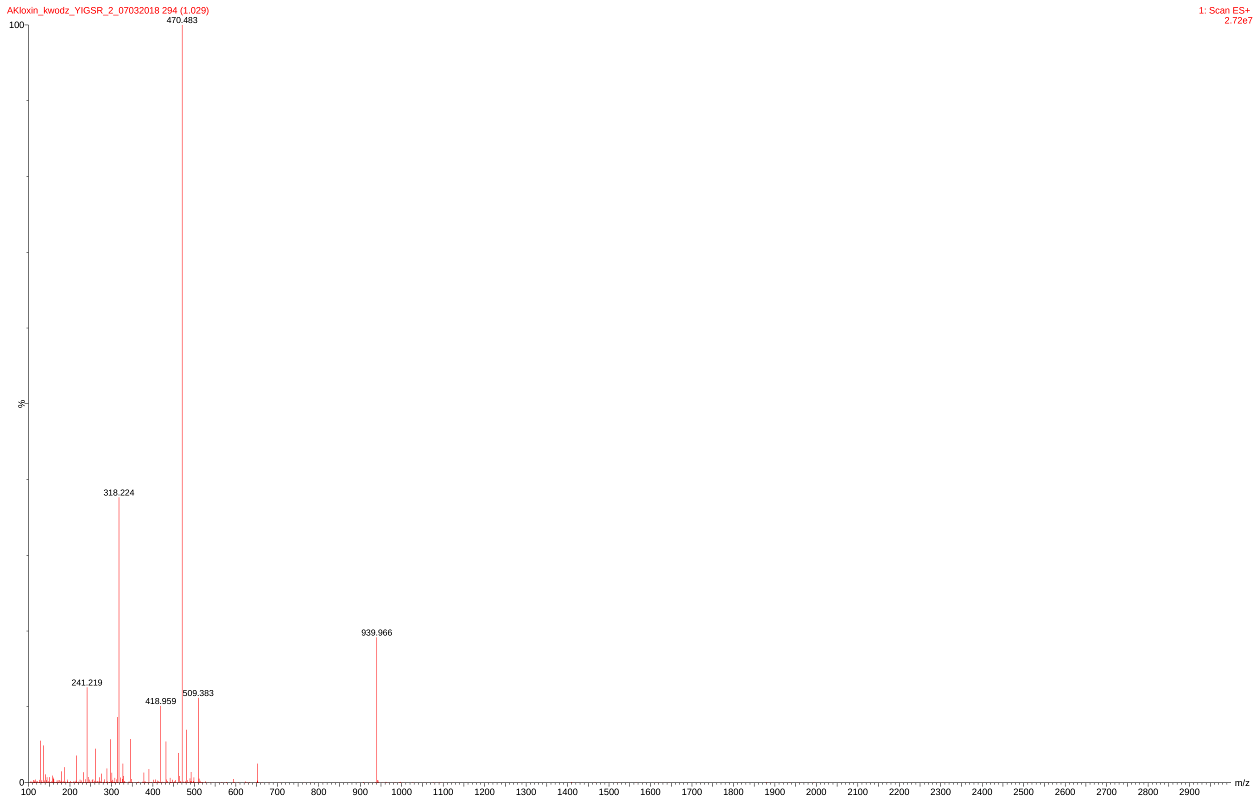


**SI Figure 6:** Mass spectrometry of monofunctional pendant peptide. Successful synthesis of pendant peptide, CGKGYIGSR-amide, was confirmed by SQD2 mass spectrometry. Expected molecular weight of 939 g/mol. $[M + H]^{+}$= 940 g/mol. $[M + 2H]^{+}$= 470 g/mol.

#### XI. NMR Spectra of PEG-8-Nb

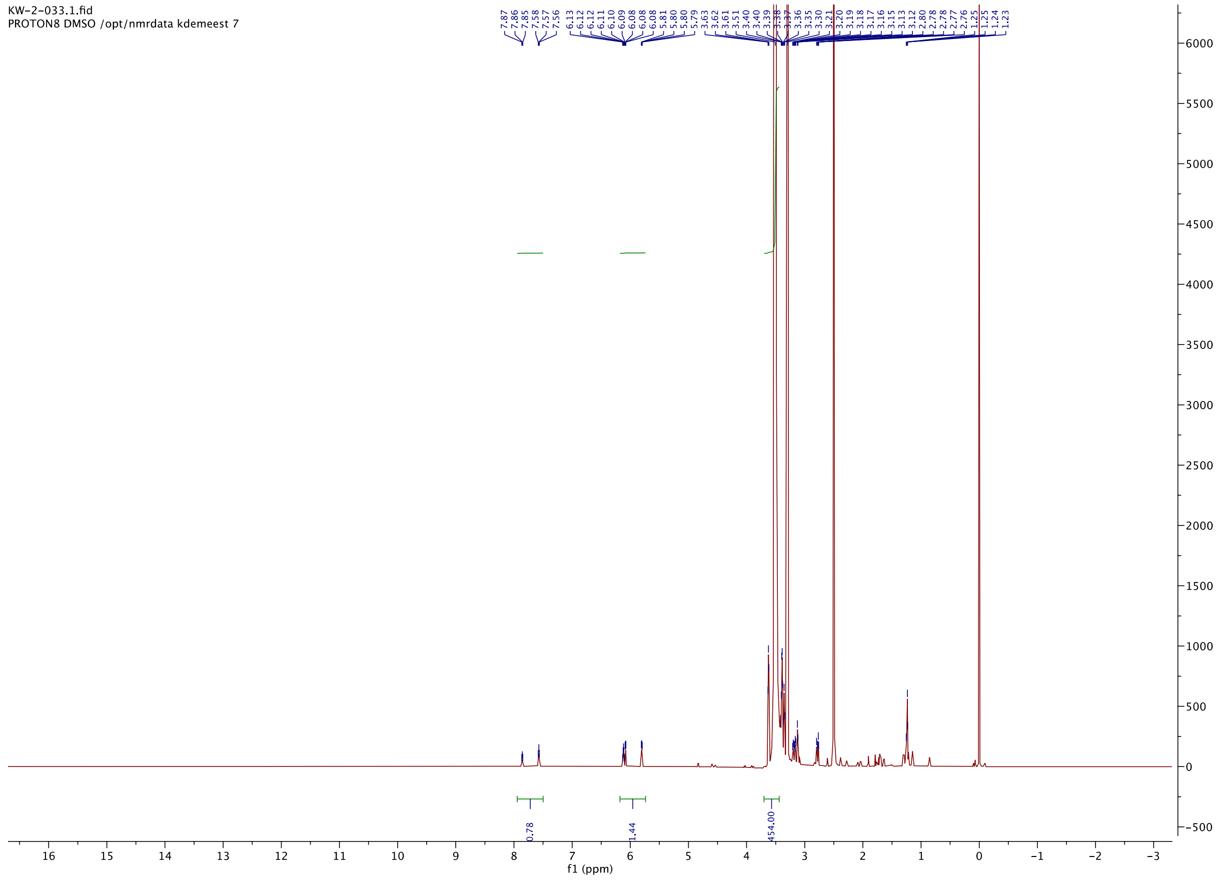


**SI Figure 7:** Representative $^1$H NMR of PEG-8-Nb. The functionality is based on the number of protons corresponding to norbornene normalized to the PEG backbone. With expected integration for norbornene protons (2H, 6.20 to 5.86 ppm) and the PEG backbone protons (454 H, 3.65 to 3.40 ppm), the calculated norbornene functionality was 1.44 divided by 2, which equals 72% functionality for this batch where ~ 75% functionality was observed on average between batches.
